## Supplementary figures and tables for "Atlas-scale single-cell multi-sample multi-condition data integration using scMerge2"

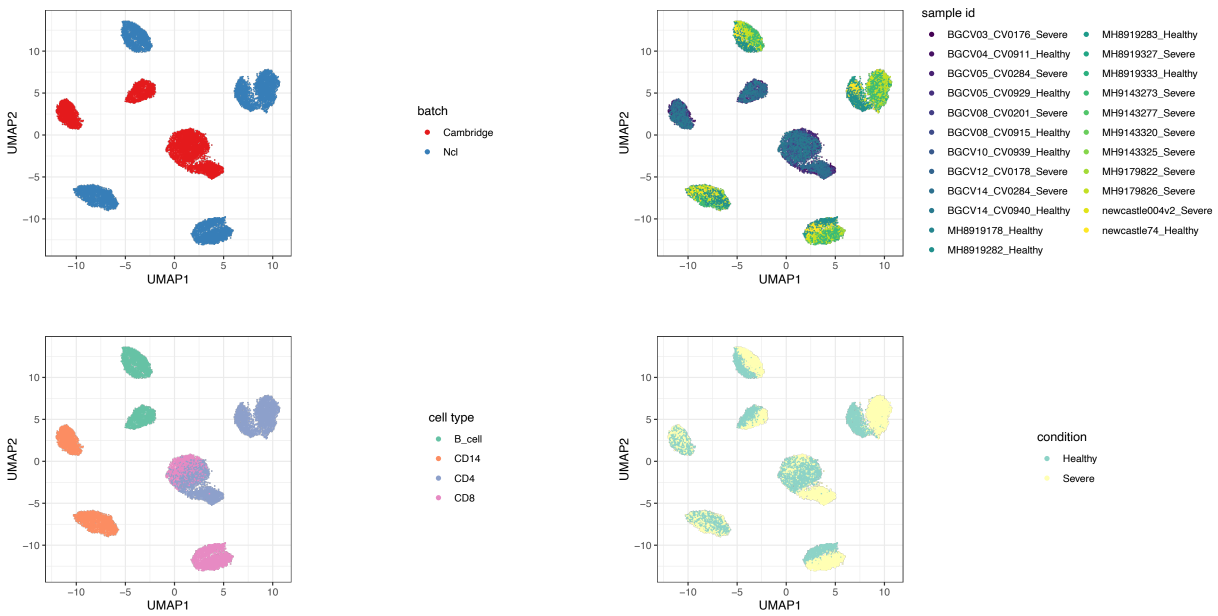


Supplementary Figure S1: UMAP plots of an example of simulated data (logFC = 1.2, DS% = 5%), coloured by batch, sample id, cell type and condition.

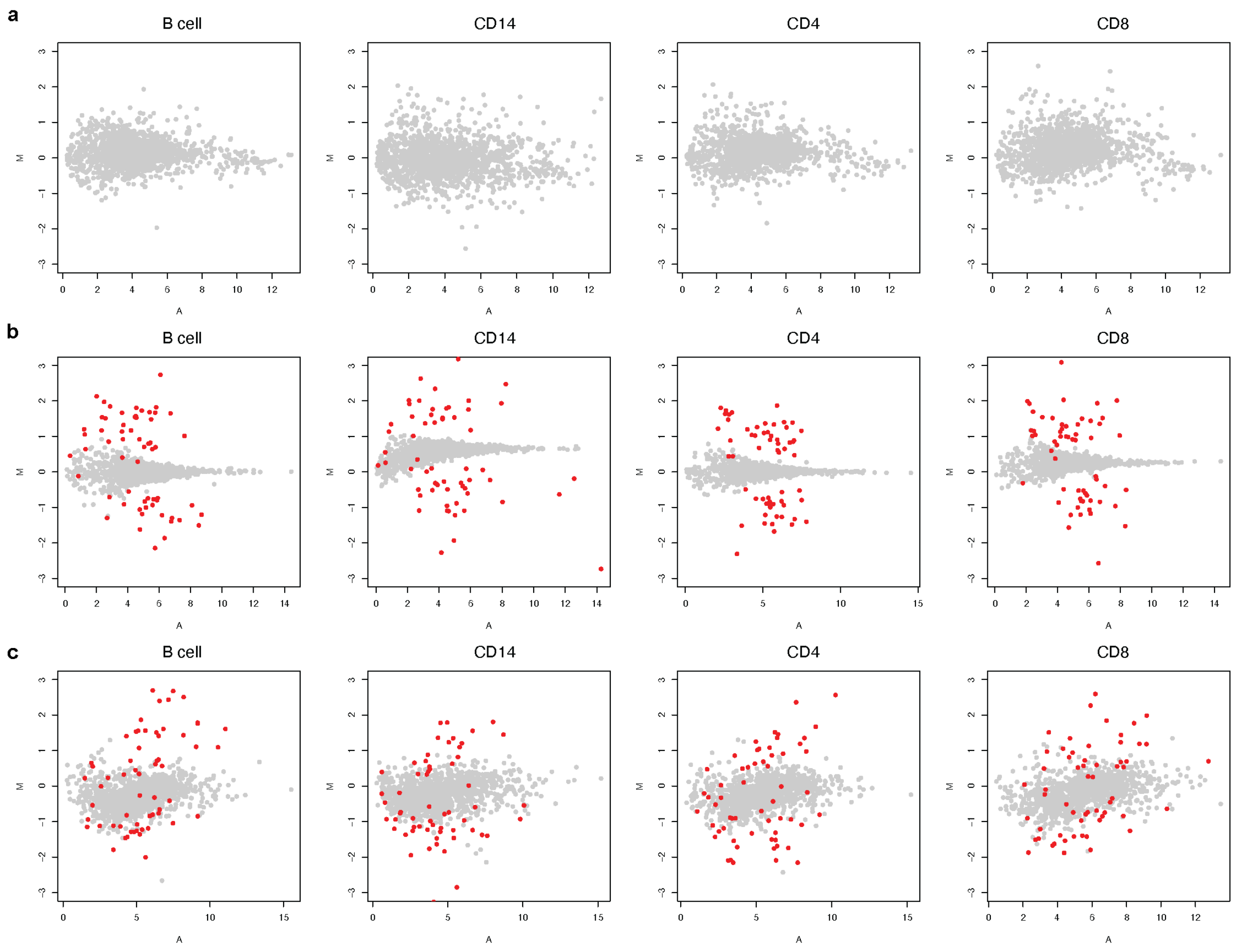


Supplementary Figure S2: MA plots of the real and simulated data, where x-axis is the average of gene expression and y-axis is the difference of the gene expression between two condition: (a) Real data; (b) Simulated data using mu formula ∼`cell type`, estimated from data with one condition; (c) Simulated data using mu formula ∼`cell type + sample ID + condition`, estimated from data from two conditions but with condition label permuted. The red dots indicates the simulated ground truth DS genes. The simulation strategy (c) exhibits a more similar pattern with the real data, which therefore is used in this study.

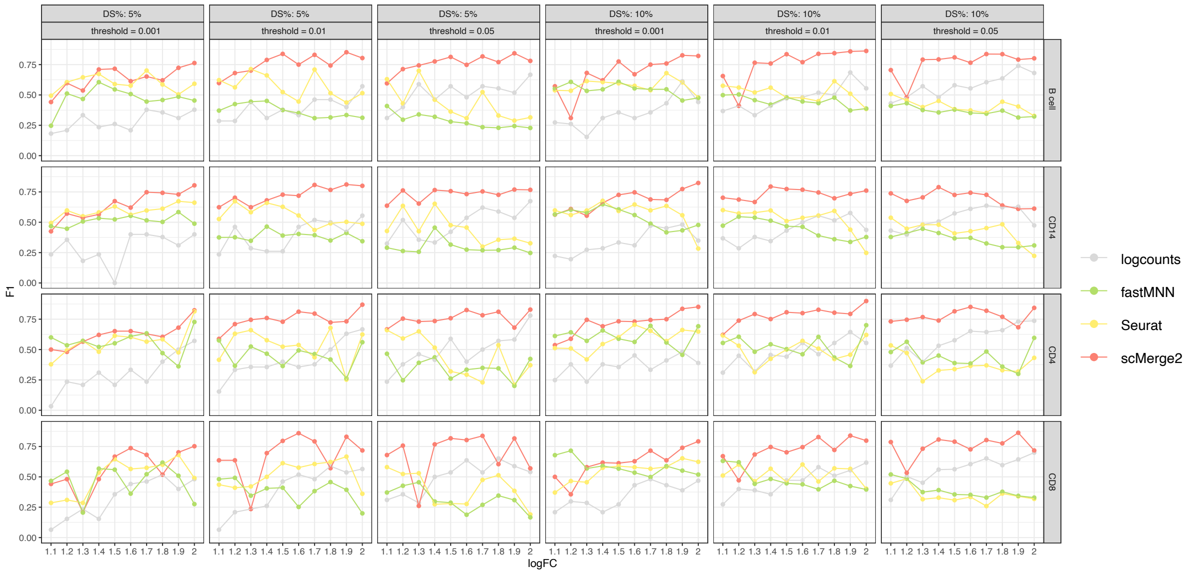


Supplementary Figure S3: F1-score of the differential state (DS) results of four cell types (B cell, CD14, CD4 and CD8) (row) of simulated data, with 5% (1st - 3rd column) and 10% DS genes (4th - 6th column) within each cell type, for scMerge2, Seurat, fastMNN and raw, varying simulated log fold change (logFC) of DS genes (x-axis) and different threshold of adjusted p-value (column).

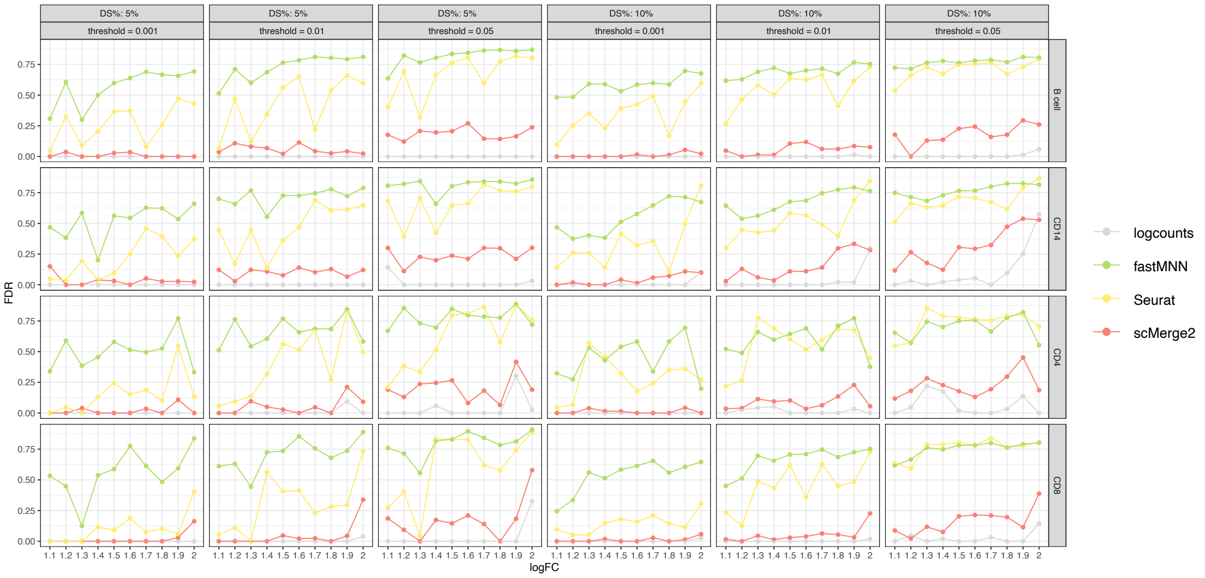


Supplementary Figure S4: FDR of the differential state (DS) results of four cell types (B cell, CD14, CD4 and CD8) (row) of simulated data, with 5% (1st - 3rd column) and 10% DS genes (4th - 6th column) within each cell type, for scMerge2, Seurat, fastMNN and raw, varying simulated log fold change (logFC) of DS genes (x-axis) and different threshold of adjusted p-value (column).

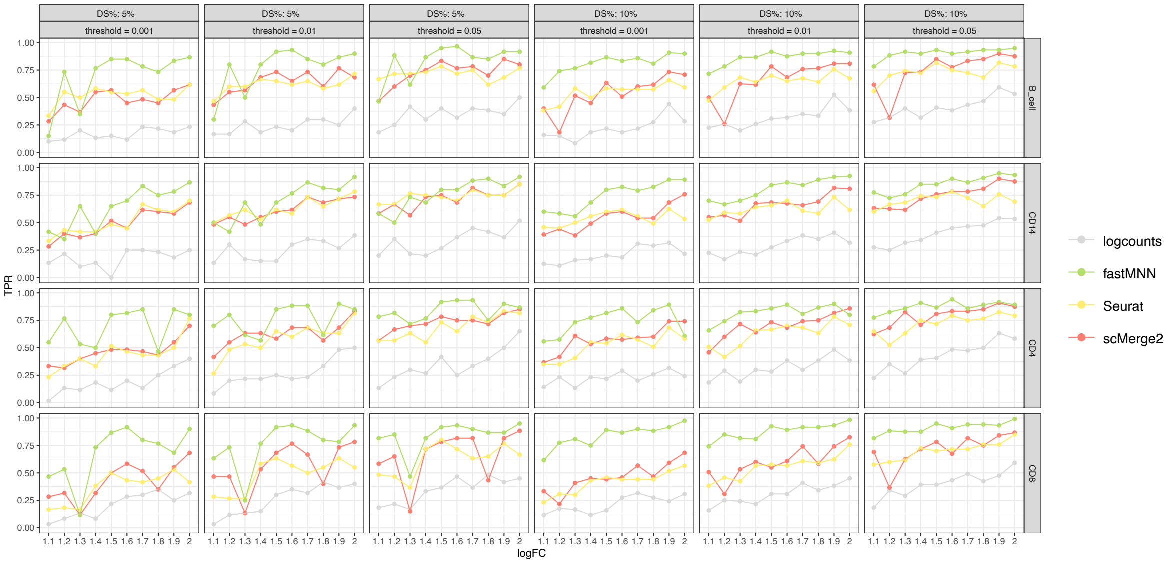


Supplementary Figure S5: TPR of the differential state (DS) results of four cell types (B cell, CD14, CD4 and CD8) (row) of simulated data, with 5% (1st - 3rd column) and 10% DS genes (4th - 6th column) within each cell type, for scMerge2, Seurat, fastMNN and raw, varying simulated log fold change (logFC) of DS genes (x-axis) and different threshold of adjusted p-value (column).

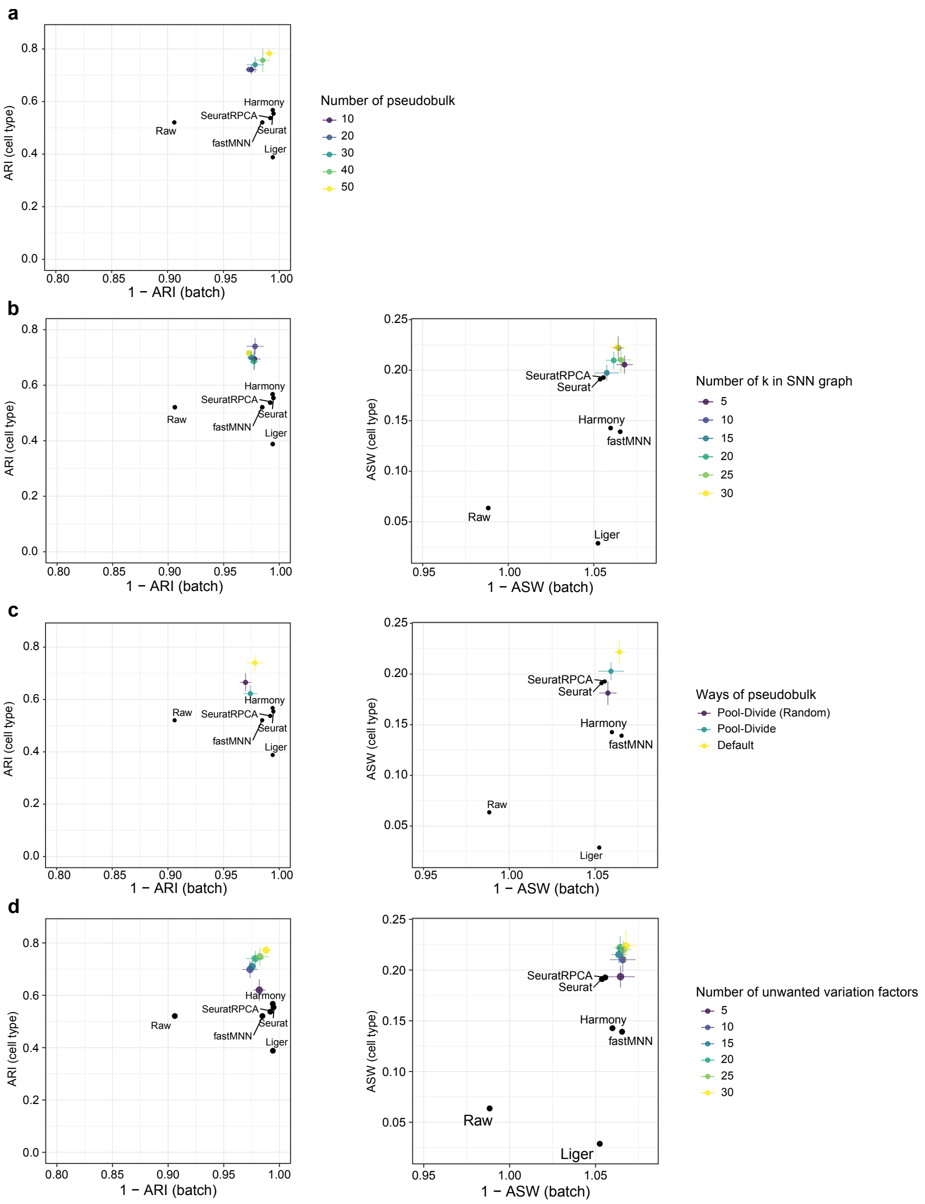


Supplementary Figure S6: Robustness analysis of the tuning parameters of scMerge2 using COVID-19 60k data: Adjusted rand index (ARI) (left panel), where x-axis indicates 1 minus batch ARI and y-axis indicates cell type ARI; Average silhouette width (ASW), where x-axis indicates 1 minus batch ASW and y-axis indicates cell type ASW (right panel), when varying (a) the number of pseudobulk constructed (10, 20, 30 (default), 40, 50); (b) the number of k used in SNN graph (5, 10 (default), 15, 20, 25, 30); (c) different methods to construct pseudobulk. (d) Number of unwatned variation factors (5, 10, 15, 20 (default), 25, 30).

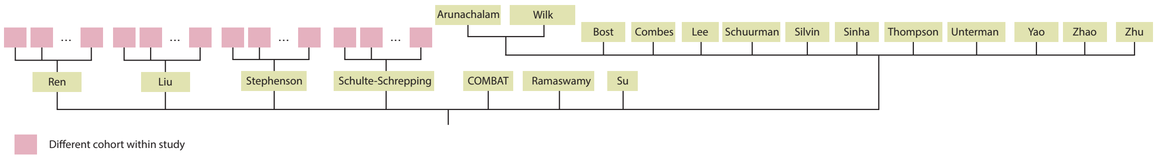


Supplementary Figure S7: Hierarchical merging strategy for COVID-19 scRNA-seq data collection.

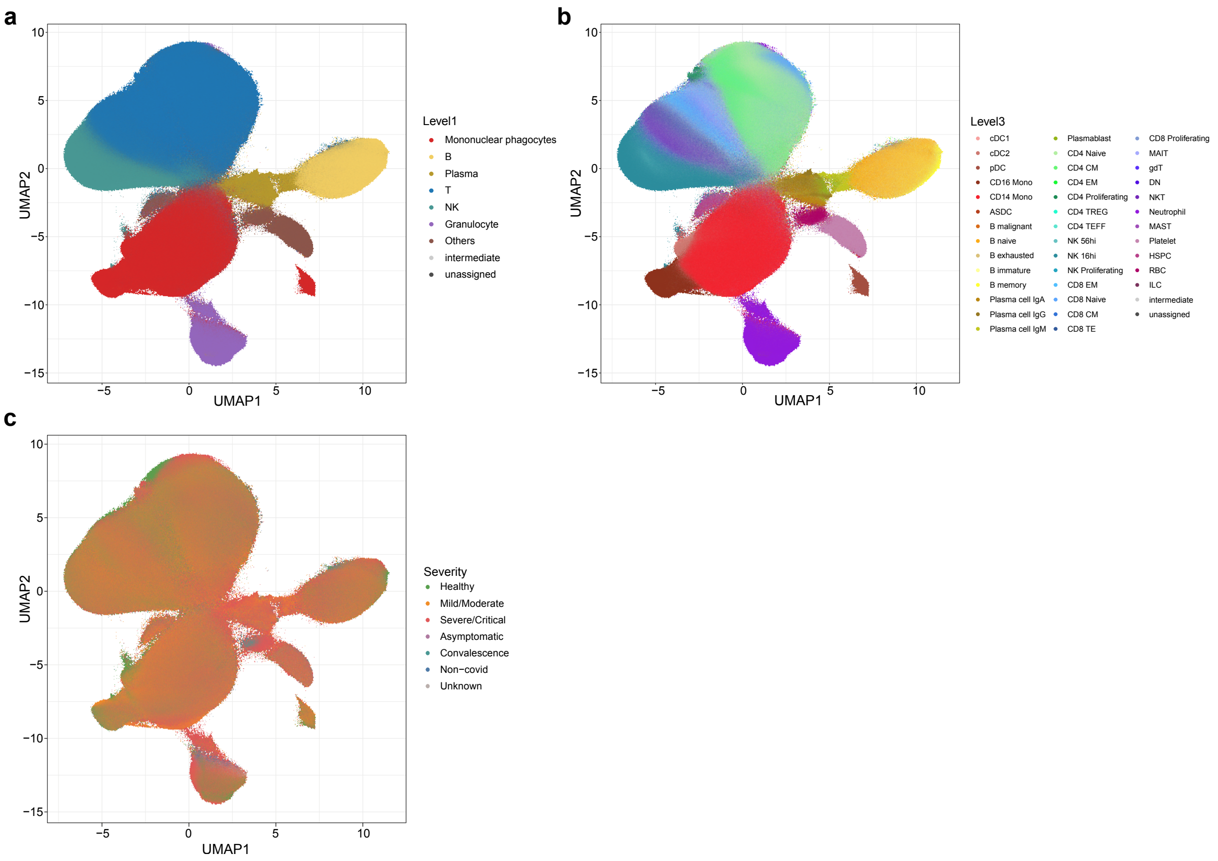


Supplementary Figure S8: UMAP of integration of COVID-19 data collection after scMerge2 integration, coloured by (a) level 1 cell type annotation; (b) level 3 cell type annotation and (c) severity.

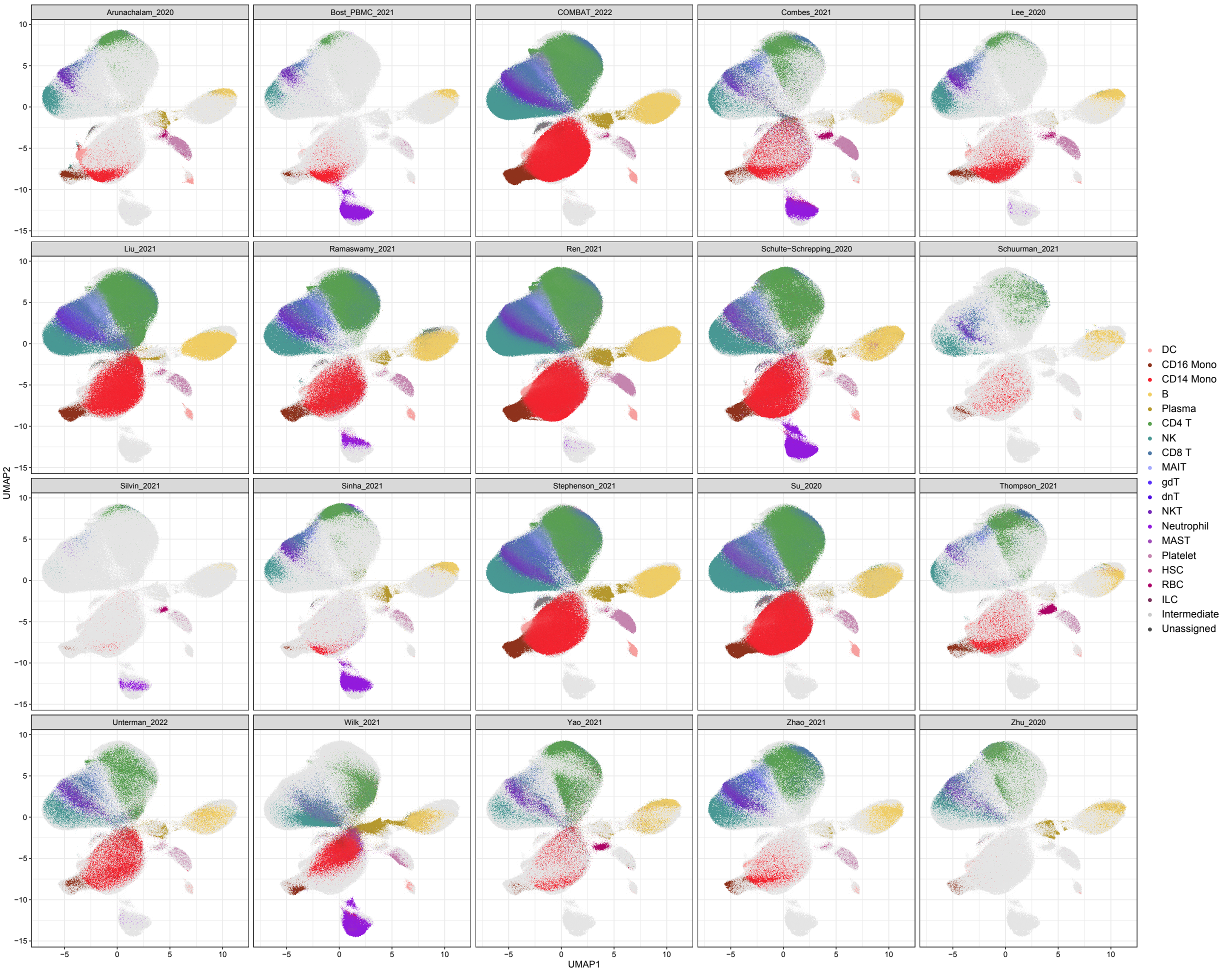


Supplementary Figure S9: UMAP of integration of COVID-19 data collection after scMerge2 integration, coloured by cell type (level 2) and faceted by dataset.

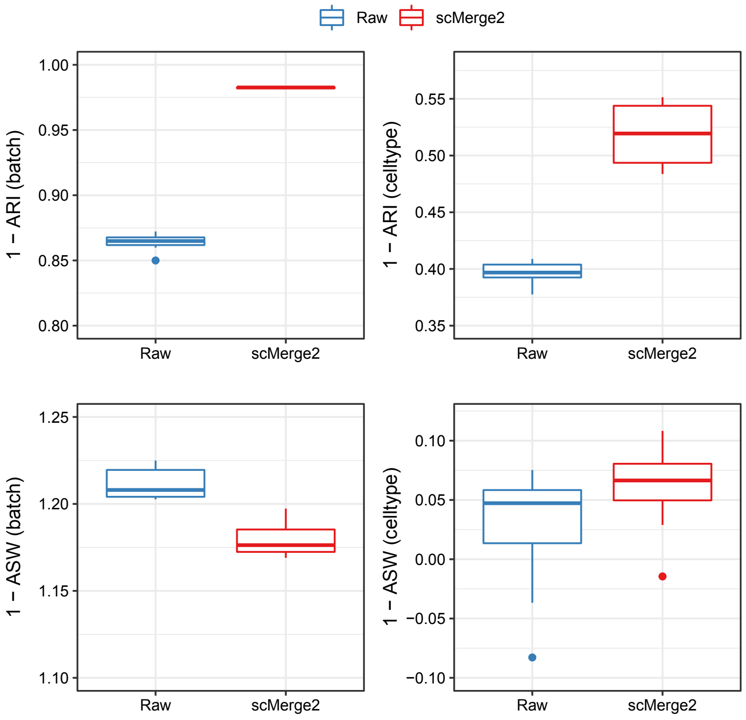


Supplementary Figure S10: Boxplots of evaluation metrics of COVID-19 scRNA-seq data collection for scMerge2-h (data merged in a hierarchical manner) and Raw, where the first row indicates the results of adjusted rand index (ARI): 1 minus batch ARI (left) and cell type ARI (right); the second row indicates the results of Average silhouette width (ASW): 1 minus batch ASW (left) and cell type ASW (right). For all of the four metrics, higher value indicates better performance. Since the size of this data collection is large, we subsampled 1% of the cells to calculate the metrics, and repeated this procedure 10 times.

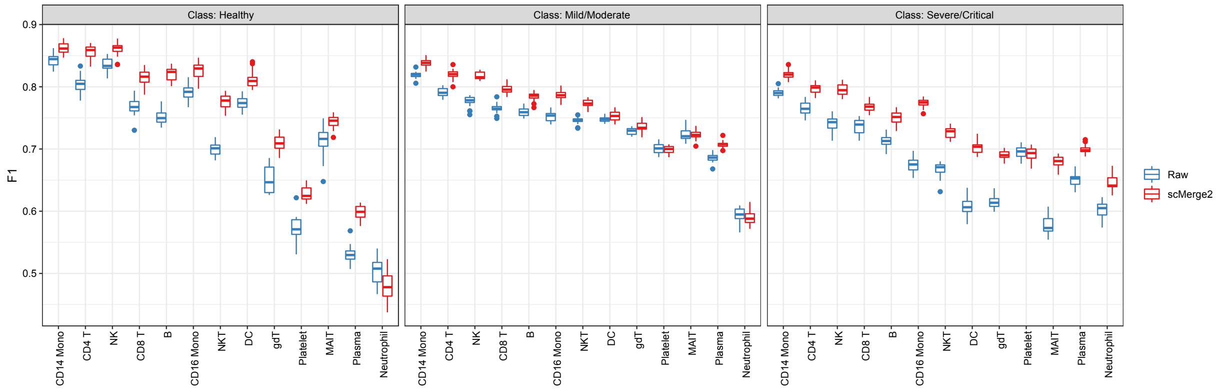


Supplementary Figure S11: Prediction results of disease severity using cell type-specific aggregated expression calculated from raw logcounts (blue) and scMerge2 adjusted results (red), evaluated by class-specific F1 scores.

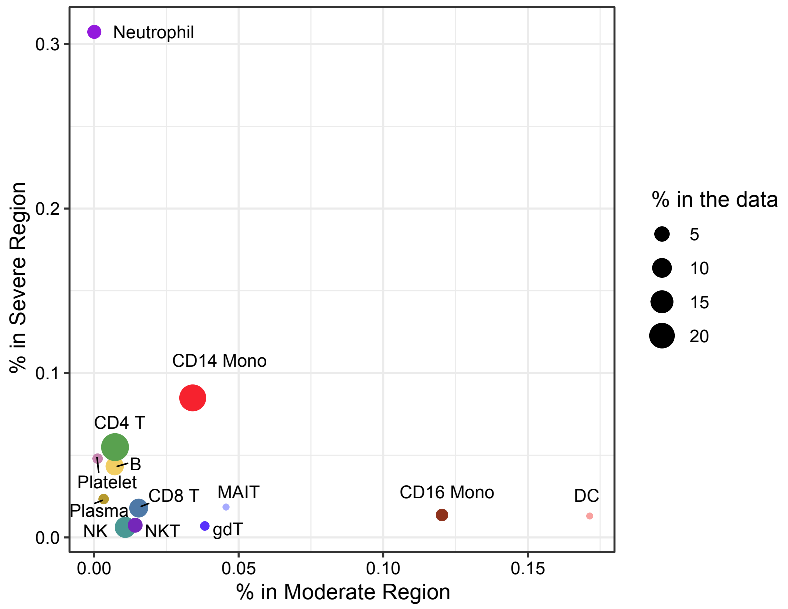


Supplementary Figure S12: Scatter plot shows the proportion of cells in Moderate region (x-axis) vs the proportion of cells in Severe region, determined by DAseq. The size of each point indicates the cell type proportion in the all data (Only cell types that have more than 1% in the data are shown).

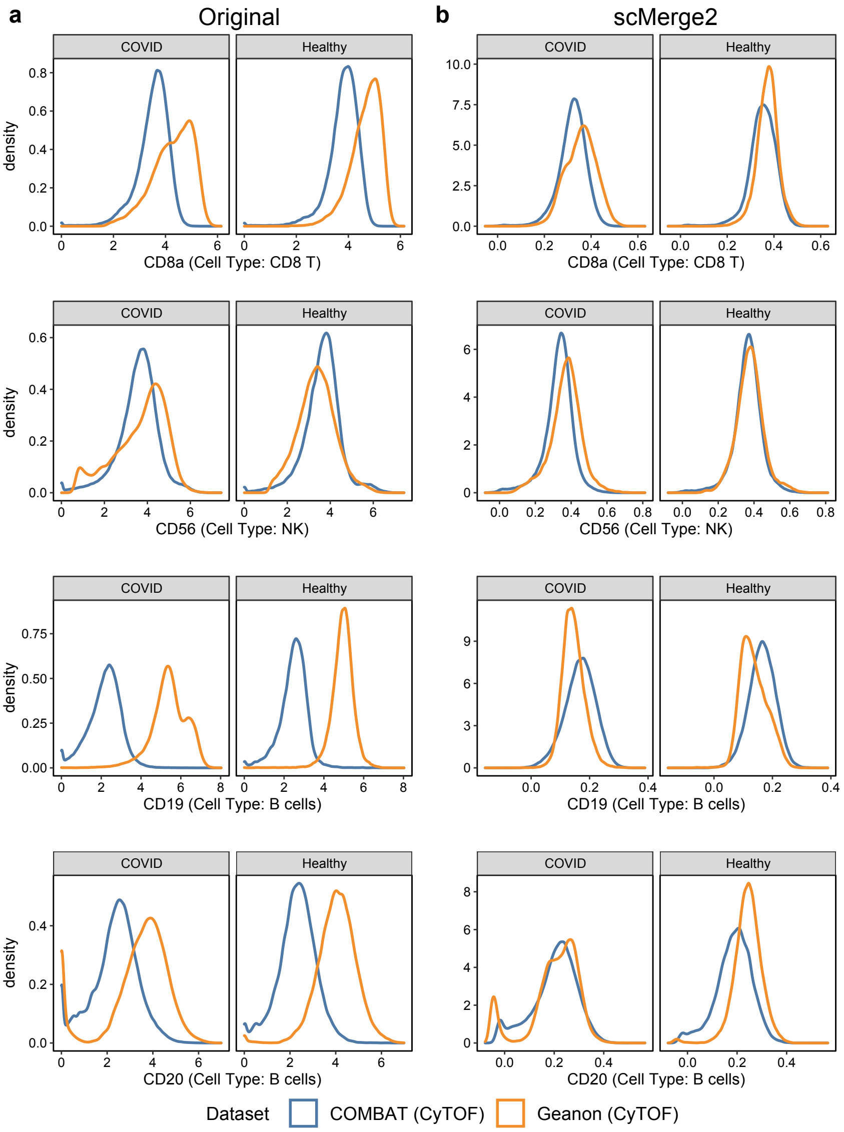


Supplementary Figure S13: Density plot of selected marker in specific cell type: CD8a in CD8 T cells; CD56 in NK cells; CD19 in B cells and CD20 in B cells, using (a) original expression and (b) scMerge2 adjusted expression. Within a specific cell type, the distribution of the cell type marker is expected to be similar between two datasets.

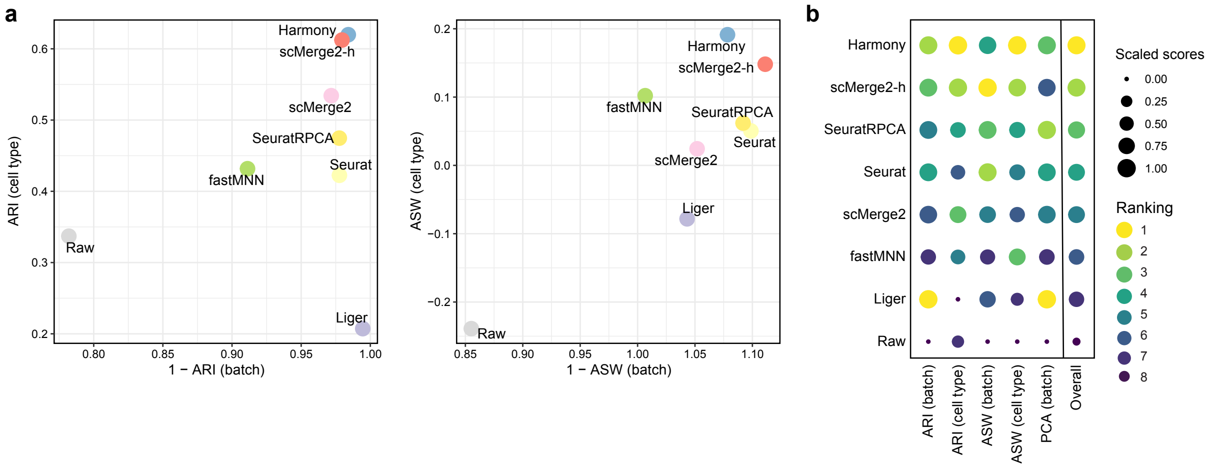


Supplementary Figure S14: CITE-seq data example: (a) Scatter plots of evaluation metrics of ADT data integration of a 200k cells subset of two COVID-19 studies (Liu and Stephenson) for scMerge2, scMerge2-h (data merged in a hierarchical manner), Seurat, Seurat (RPCA), Harmony, fastMNN, Liger and Raw: Adjusted rand index (ARI) (left panel), where x-axis indicates 1 minus batch ARI and y-axis indicates cell type ARI; Average silhouette width (ASW), where x-axis indicates 1 minus batch ASW and y-axis indicates cell type ASW (right panel). (b) Dot plots indicates the ranking of the data integration methods in terms of 5 different evaluation metrics. The size of the dot indicates the scaled scores, which are obtained from the min-max scaling of the original values. The overall ranking is ranked based on the average ranking of the five evaluation metrics.

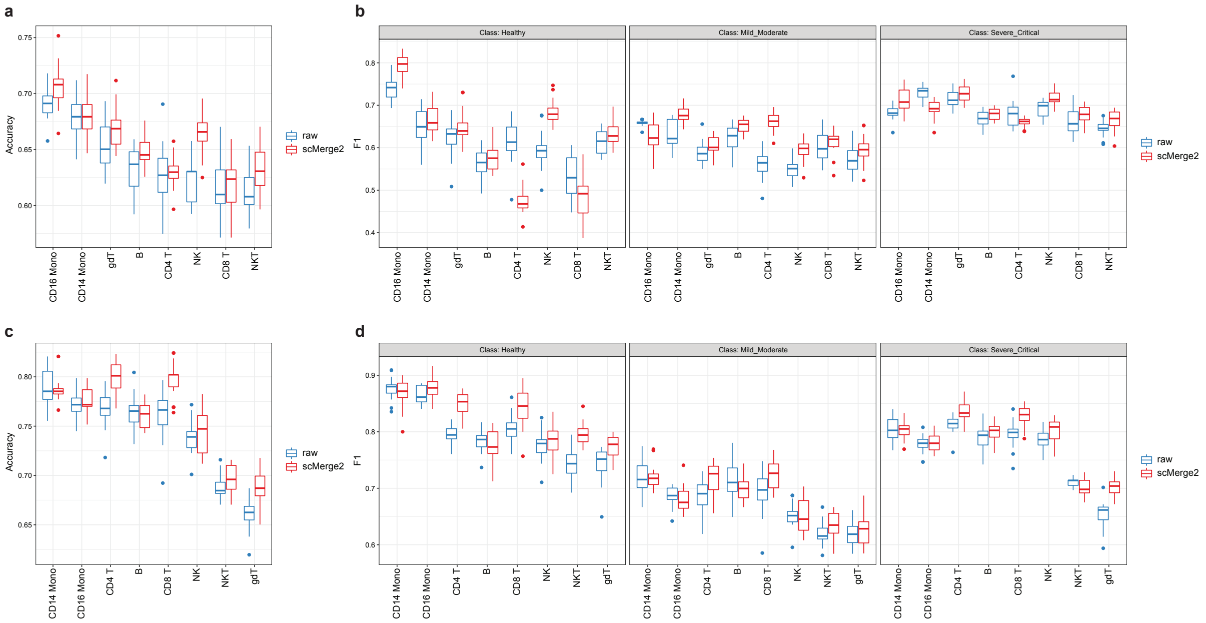


Supplementary Figure S15: CITE-seq data example: Prediction results of disease severity using cell type-specific aggregated expression calculated from raw logcounts (blue) and scMerge2 normalised results (red), using (a-b) ADT expression and (c-d) RNA expression.

**Supplementary Table 1**. Data collections used in the paper.

| | Study | Accession | Number of cells | Number of samples | Number of donors | Country | Published Date | doi |
|---|---|---|---|---|---|---|---|---|
| | Arunachalam_2020 | GSE155673 | 56639 | 12 | 12 | US | August 11, 2020 | https://www.science.org/doi/10.1126/science.abc6261 |
| | Bost_PBMC_2021 | GSE157344 | 50284 | 33 | 33 | Isreal | March 5, 2021 | https://doi.org/10.1038/s41467-021-21702-6 |
| | COMBAT_2022 | EGAS00001005493 | 783704 | 140 | 140 | UK | March 3, 2022 | https://doi.org/10.1016/j.cell.2022.01.012 |
| | Combes_2021 | GSE163668 | 111990 | 44 | 44 | US | January 25, 2021 | https://doi.org/10.1038/s41586-021-03234-7 |
| | Lee_2020 | GSE147507 | 59572 | 20 | 17 | Korea | July 10, 2020 | https://www.science.org/doi/10.1126/sciimmunol.abd15544 |
| | Liu_2021 | GSE161918 | 411902 | 70 | 47 | US | April 1, 2021 | https://doi.org/10.1016/j.cell.2021.02.0188 |
| | Ramaswamy_2021 | GSE166489 | 271267 | 38 | 32 | US | May 11, 2021 | https://doi.org/10.1016/j.immuni.2021.04.003 |
| | Ren_2021 | GSE158055 | 999462 | 173 | 151 | China | April 1, 2021 | https://doi.org/10.1016/j.cell.2021.01.053 |
| | Schulte-Schrepping_2020 | EGAS00001004571 | 328780 | 147 | 74 | Germany | September 17, 2020 | https://doi.org/10.1016/j.cell.2020.08.001 |
| **COVID-19 scRNA-seq** | Schuurman_2021 | GSE164948 | 32384 | 20 | 20 | Netherlands | August 23, 2021 | https://doi.org/10.7554/eLife.696611 |
| | Silvin_2020 | E-MTAB-9221 | 6960 | 10 | 10 | France | September 17, 2020 | https://doi.org/10.1016/j.cell.2020.08.002 |
| | Sinha_2022 | GSE157789 | 80994 | 21 | 14 | Canada | January 1, 2022 | https://doi.org/10.1038/s41591-021-01576-3 |
| | Stephenson_2021 | E-MTAB-10026 | 643071 | 143 | 130 | UK | April 20, 2021 | https://doi.org/10.1038/s41591-021-01329-2 |
| | Su_2020 | E-MTAB-9357 | 538210 | 268 | 143 | US | October 10, 2020 | https://doi.org/10.1016/j.cell.2020.10.037 |
| | Thompson_2021 | GSE166992 | 63895 | 8 | 8 | US | March 16, 2021 | https://doi.org/10.1016/j.celrep.2021.108863 |
| | Unterman_2022 | GSE155224 | 80789 | 18 | 10 | US | January 22, 2022 | https://doi.org/10.1038/s41467-021-27716-4 |
| | Wilk_2021 | GSE174072 | 174753 | 55 | 39 | US | June 15, 2021 | https://doi.org/10.1084/jem.20210582 |
| | Yao_2021 | GSE154567 | 69983 | 17 | 17 | US | January 5, 2021 | https://doi.org/10.1016/j.celrep.2020.108590 |
| | Zhao_2021 | CNP0001250 | 88374 | 38 | 19 | China | September 16, 2021 | https://doi.org/10.1038/s41392-021-00753-7 |
| | Zhu_2020 | CNP0001102 | 46022 | 23 | 3 | China | September 15, 2020 | https://doi.org/10.1016/j.immuni.2020.07.009 |
| | | Total: | 4899035 | 1298 | 963 | | | |
| **COVID-19 CyTOF** | COMBAT_2022 | EGAS00001005493 | 7118158 | 160 | 160 | UK | March 3, 2022 | https://doi.org/10.1016/j.cell.2022.01.012 |
| | Geanon_2022 | FR-FCM-Z2XA | 4747543 | 21 | 21 | US | February 16, 2021 | https://doi.org/10.1002/cyto.a.24317 |
| | | Total: | 11865701 | 181 | 181 | | | |

| | | | | | | | | |
|---|---|---|---|---|---|---|---|---|
| **COVID-19 200k CITE-seq (COVID-19 200k)** | Liu_2021 | GSE161918 | 82537 | 70 | 47 | US | April 1, 2021 | https://doi.org/10.1016/j.cell.2021.02.0188 |
| | Stephenson_2021 | E-MTAB-10026 | 117463 | 114 | 104 | UK | April 20, 2021 | https://doi.org/10.1038/s41591-021-01329-2 |
| | | Total: | 200000 | 184 | 151 | | | |
| **COVID-19 60k** | Stephenson_2021 | E-MTAB-10026 | 66967 | 58 | 53 | UK | April 20, 2021 | https://doi.org/10.1038/s41591-021-01329-2 |
| **COVID IMC** | Rendeiro_2021 | zenodo.4110560, zenodo.4139442, zenodo.4637034 | 664006 | 237 | 23 | US | March 29, 2021 | https://doi.org/10.1038/s41586-021-03475-6 |
